# HIR-1 Mediates Response to Hypoxia-Induced Extracellular Matrix Remodeling

**DOI:** 10.1101/302638

**Authors:** Roman Vozdek, Yong Long, Dengke K. Ma

## Abstract

Inadequate tissue oxygen, or hypoxia, is a central concept in pathophysiology of ischemic disorders and cancer. Hypoxia promotes extracellular matrix (ECM) remodeling, cellular metabolic adaptation and metastasis. To determine how cells respond to hypoxia-induced ECM remodeling, we performed a large-scale forward genetic screen in *C*. *elegans*. We identified a previously uncharacterized receptor tyrosine kinase (RTK) named HIR-1 as a key mediator in a pathway that orchestrates transcriptional responses to hypoxia-induced ECM remodeling. Impaired ECM integrity caused by hypoxia or deficiency of the oxygen-dependent procollagen hydroxylases, heme peroxidases or cuticular collagens activates gene expression through inhibition of HIR-1. Genetic suppressor screens identified NHR-49 and MDT-15 as transcriptional regulators downstream of HIR-1. Cellular responses through HIR-1 maintain ECM homeostasis and promote animal adaptation to severe hypoxia. We propose that *C*. *elegans* HIR-1 defines an unprecedented type of RTK that mediates responses to hypoxia-induced ECM remodeling by mechanisms that are likely conserved in other organisms.

**ONE-SENTENCE SUMMARY:** A regulatory pathway for ECM homeostasis underlies adaptation to hypoxia and re-oxygenation

## INTRODUCTION

Oxygen is essential for aerobic metabolism of life. Varying oxygen levels occur in natural environments and in tissues of living organisms, eliciting highly orchestrated organismic and cellular responses to maintain proper metabolic and physiological homeostasis. For example, mammals adjust pulmonary ventilation and blood circulation to improve oxygen delivery to target tissues under hypoxic conditions, and the roundworm *C*. *elegans* can navigate for preferred oxygen levels across a gradient of ambient hypoxia and trigger rapid locomotor response upon severe hypoxia and restoration of oxygen (1–3). Hypoxia is also a common pathophysiological condition in human disorders characterized by a low supply of oxygen, including myocardial ischemia, stroke and tumorigenesis (4). Pathological hypoxic conditions can lead to tissue necrosis and fibrosis, degeneration, inflammation and tumor metastasis, driving disease progression and leading to organismic mortality (5).

Animals respond to chronic hypoxia through evolutionarily conserved molecular pathways and cellular mechanisms that regulate gene expression and reprogram metabolism (4,6,7). The hypoxia inducible factor (HIF) is a master transcriptional regulator of hypoxic responses. Genetic studies of *C*. *elegans* led to the discovery of the evolutionarily conserved family of HIF hydroxylases (EGL-9 in *C*. *elegans* and EGLN2 in humans) that link oxygen-sensing to HIF-1 activation and transcriptional responses to hypoxia (8,9). Hydroxylated HIF under normoxia is recognized by the Von Hippel-Lindau tumor suppressor and targeted for proteasomal degradation whereas hypoxia causes impaired HIF hydroxylation, leading to transcriptional activation of HIF target genes (10). In *C*. *elegans*, the HIF-1 pathway mediates various physiological and behavioral responses to hypoxia (2,6,11,12). The transcriptional targets of mammalian HIF include *LOX*, *LOXL2* and *LOXL4* encoding copper-dependent lysyl oxidases that promote cross-linking of ECM components for enhanced tissue stiffness, a proposed trigger of tumor metastasis (5). Hypoxia-induced ECM remodeling, in turn, activates intracellular signaling cascades to regulate cell fate, metastasis and adaptation to hypoxia (13,14). How cells sense and respond to hypoxia-induced ECM remodeling remains undefined.

Collagens are major integral components of ECM that are extensively modified by oxygen-dependent enzymes including lysyl oxidases, lysyl hydroxylases and prolyl hydroxylases. Beyond HIF dependent mechanisms, it is unknown whether hypoxia can alter ECM integrity directly by impairing oxygen-dependent collagen modification and how ECM remodeling might elicit subsequent HIF-independent gene regulation. In mammalian cells, *VEGF* (vascular endothelial growth factor) can be induced by hypoxia through both HIF-dependent and HIF-independent mechanisms (15). Various protein kinases and transcription factors other than HIF respond to hypoxia in a HIF-independent manner to modulate transcriptional responses (16,17). Mitogen activated protein kinases or integrin-linked protein kinases can transduce remodeled ECM signals to influence cell fate and resistance to hypoxia through transcriptional regulation (13,14). *C*. *elegans* has also been extensively used to investigate HIF-independent responses that mediate physiological and behavioral adaptation to severe hypoxia (18–28). Nonetheless, the precise roles and cellular mechanisms of HIF-independent transcriptional response to hypoxia-induced ECM remodeling in physiology and diseases remain poorly understood.

We sought to identify new genes and pathways that transduce hypoxic signals to gene regulation independently of HIF-1. We generated *C*. *elegans* transgenic animals carrying GFP reporters that were robustly activated by hypoxia even in HIF-1-deficient animals. We performed large-scale forward genetic screens to isolate mutants with constitutively activated reporters under normoxia and performed suppressor screens to identify hypoxia-regulated transcription factors. We found that exposure to hypoxia resulted in the remodeling of ECM, which in turn triggers intracellular transcriptional responses by inhibiting a cell transmembrane RTK that we named HIR-1 (<u>H</u>ypoxia <u>I</u>nhibited <u>R</u>eceptor tyrosine kinase).

## RESULTS

### *comt*-*5p::GFP* is a robust live reporter induced by hypoxia independently of HIF-1

To identify genes and pathways that mediate HIF-1-independent transcriptional response by hypoxia, we used RNA-seq to compare transcriptomes of wild-type (WT) *C*. *elegans* under normoxia (21% oxygen) or severe hypoxia (nearly 0% oxygen) for 2 hours, and *egl*-*9* null mutants (in which HIF-1 is constitutively activated) under normoxia. We identified 72 genes that showed increased expression after hypoxia in WT animals but not in *egl*-*9* null mutants (Supplementary Fig. S1A). We generated transgenic animals with green fluorescent protein (GFP) driven by the promoters of these genes and focused on one of the reporters constructed, *dmaIs1*, that showed robust induction of GFP in the hypodermal and intestinal cells within 24 hours in 0.5% O_2_ (Fig. 1A and 1B). *dmaIs1* carries GFP driven by the promoter of *comt*-*5*, which encodes a predicted catechol-*O*-methyltransferase (Fig. 1C). We crossed animals carrying the *dmaIs1* transgene with *egl*-*9*(*sa307*) or *hif*-*1*(*ia04*) loss-of-function (LOF) deletion mutants and, as expected, we did not observe activation of *comt*-*5p::GFP* in *egl*-*9* mutants, whereas both WT and *hif*-*1* mutants exhibited increased expression of *comt*-*5p::GFP* by hypoxia (Fig. 1A and 1D).

**Fig. 1.**
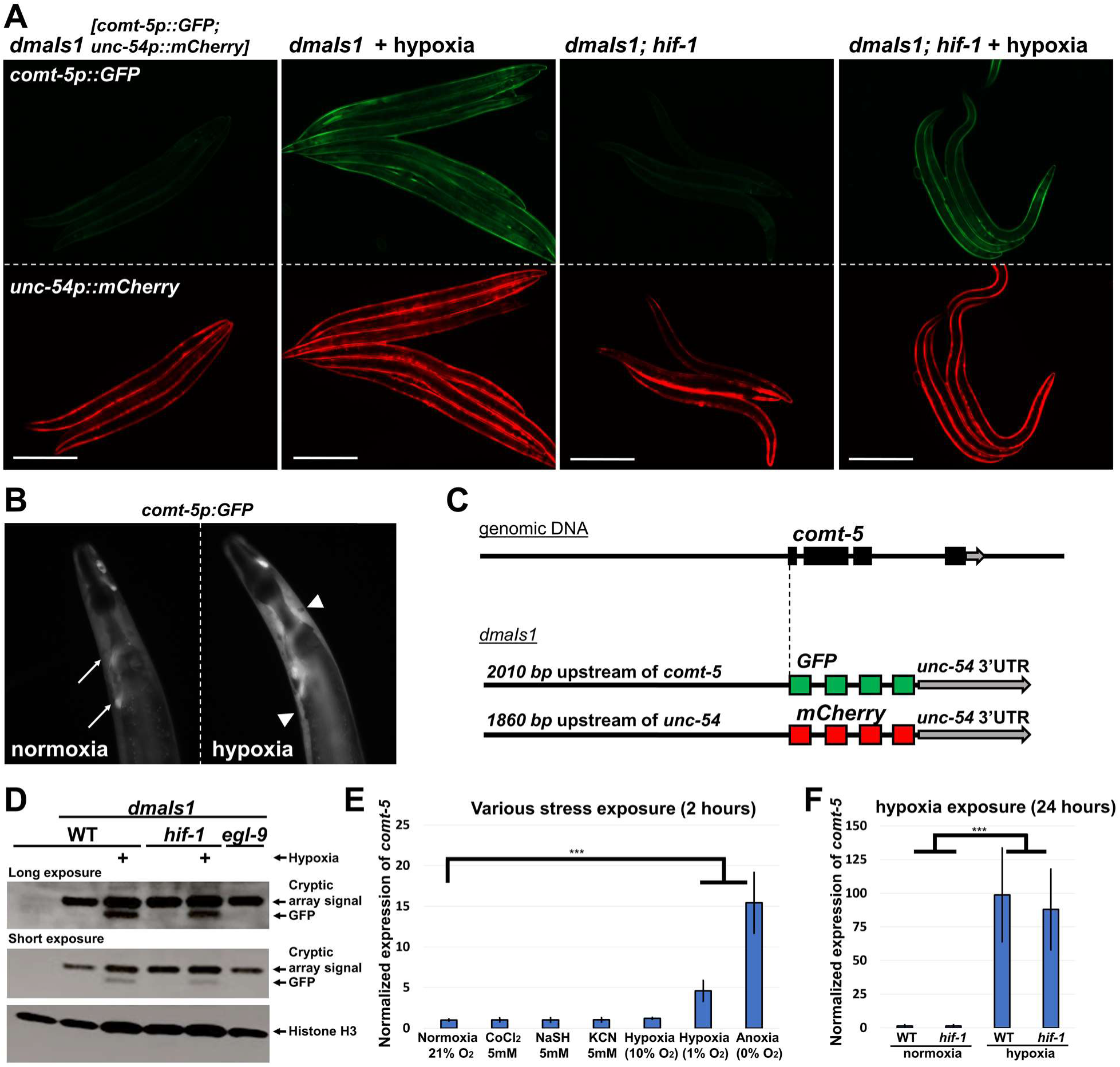
*comt*-*5p::GFP* is upregulated by hypoxia in a *hif*-*1* -independent manner. **A)** Fluorescent images of WT and *hif*-*1(ia04)* mutants carrying the genome-integrated *dmaIs1 [comt*-*5p::GFP]* transgene, exposed to normoxia or hypoxia. Representative of >100 animals **B)** Enlarged gray-scale images of *C*. *elegans* carrying *comt*-*5p::GFP* under normoxia and hypoxia. Arrows in normoxia image indicate head neurons with GFP signal. Arrowheads in hypoxia image indicate hypodermal cells and intestine with GFP signal. Representative of >100 animals **C)** Schematic diagram of the *dmaIs1* transgene. 2010 *bp* of upstream sequence of the *comt*-*5* gene was cloned before the GFP-coding sequence followed by 3’UTR of the *unc*-*54*. The *dmaIs1* transgene also carries *unc*-*54p::mCherry* as an additional marker of expression. **D)** Analysis of the GFP signal by Western blot analysis of 20 randomly picked transgenic animals carrying the *dmaIs1 [comt-5p::GFP]* transgene. Histone H3 was used as a loading control. Transgenic animals express a cryptic nonspecific protein recognized by the GFP antibody (33 kDa) that does not contribute to fluorescence in living animals. Representative of 3 independent experiments. **E)** qRT-PCR analysis of *comt*-*5* expression in animals exposed to hypoxia (1% O_2_), anoxia (0% O_2_), or hypoxia-mimicking conditions (CoCl_2_, NaSH, KCN). n ≥ 200 total animals for each group with 3 independent biological replicates; ^∗∗^ indicates P < 0.001. **F)** qRT-PCR analysis of *comt*-*5* expression in normoxia- or hypoxia-exposed WT and *hif*-*1(ia04)* animals. n ≥ 200 total animals for each group with 3 independent biological replicates; ^∗∗∗^ indicates P < 0.001. Scale bars: 300 μm.

To verify that endogenous *comt*-*5* was induced by hypoxia independently of *hif*-*1*, we quantified its expression by quantitative PCR (qRT-PCR) and found that *comt*-*5* expression was increased by hypoxia in both WT and *hif*-*1(ia04)* mutants (Fig. 1F). Other stress conditions that mimic hypoxia in activating HIF-1, including exposure of animals to CoCl_2_, H2O_2_, CN^-^ or H_2_S, did not induce the *comt*-*5p::GFP* reporter or increase endogenous *comt*-*5* expression (Fig. 1E). These results identify *comt*-*5* as a specific and robust reporter gene that is activated by hypoxia independently of EGL-9 and HIF-1.

### HIR-1 is a cell-autonomous regulator of *comt*-*5* that mediates hypoxia signaling

To identify genes and pathways mediating HIF-independent regulation of *dmaIs1*, we performed large-scale forward genetic screens using ethyl methanesulfonate (EMS)-induced mutagenesis and isolated many mutants with constitutively activated *comt*-*5p::GFP* reporters even under normoxia (Table 1). Genetic linkage analysis together with whole-genome sequencing and subsequent RNA interference (RNAi) of candidate genes identified *dma51* as an allele of the previously uncharacterized gene *C24G6*.*2* (*hir*-*1*), which encodes a predicted transmembrane protein belonging to the RTK (InterPro scan) superfamily (29) (Fig. 2D). *dma51* causes a LOF nonsense mutation W551Stop in HIR-1. We used CRISPR to generate a whole-gene deletion allele *dma101* for *hir*-*1*, which also showed constitutive *comt*-*5p::GFP* induction. A similar phenotype was also observed in mutants with partial in-frame deletion (*tm3911*) or an out-of-frame deletion (*tm4098*) that covers the exons encoding the intracellular domain of HIR-1 (Fig. 2A and 2B). A transcriptional reporter *hir*-*1p::GFP* revealed ubiquitous expression of *hir*-*1* that was especially strong in the pharynx, hypodermal and seam cells in all developmental stages (Fig. 2E). To determine whether *hir*-*1* regulates *comt*-*5p::GFP* cell-autonomously in the hypoderm, where *comt*-*5p::GFP* is induced by hypoxia, we generated transgenic animals with hypodermal-specific expression of *hir*-*1(+)* driven by the *dpy*-*7* promoter. We observed that transgenic *dpy*-*7p::hir*-*1(+)* extrachromosomal arrays rescued the *hir*-*1*(LOF)-induced hypodermal activation of *comt*-*5p::GFP* (Fig. 2F). To rule out a possibility that *hir*-*1* works cell non-autonomously, we generated transgenic animals with neuronal-specific expression of *hir*-1(+) driven by the pan-neuronal promoter *ric-19p*. We did not observe rescue of the *hir*-*1* (-)- induced activation of *comt*-*5p::GFP* (Fig. S2). These results identify *hir*-*1* as a cell-autonomous negative regulator of *comt*-*5p::GFP*.

**Table 1.**
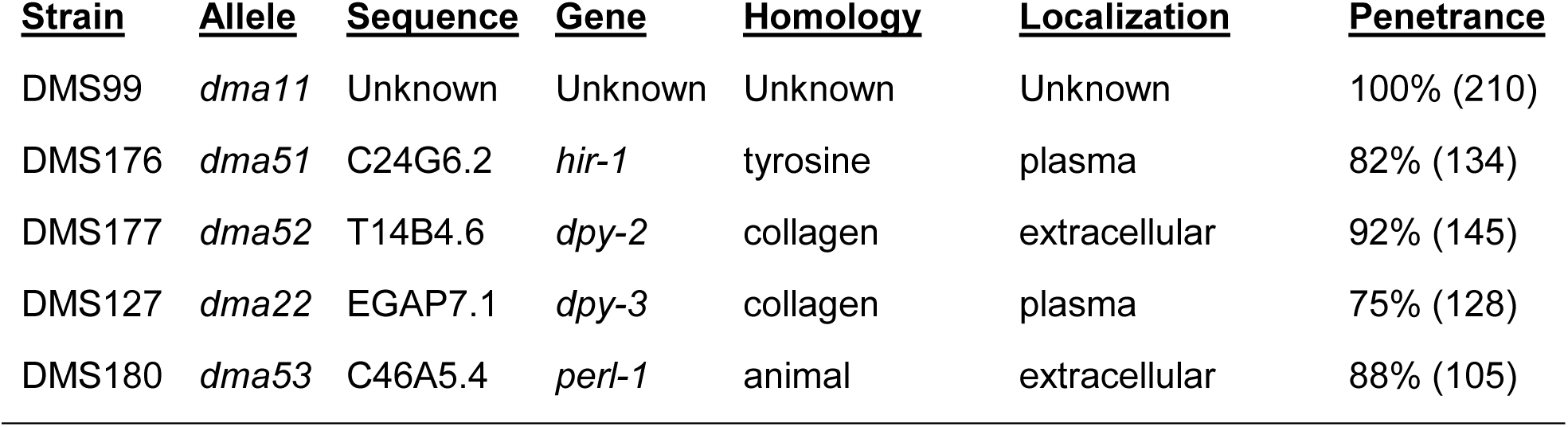
EMS-derived mutants with constitutive *comt*-*5p::GFP* expression.

| <u>Strain</u> | <u>Allele</u> | <u>Sequence</u> | <u>Gene</u> | <u>Homology</u> | <u>Localization</u> | <u>Penetrance</u> |
| --- | --- | --- | --- | --- | --- | --- |
| DMS99 | *dma11* | Unknown | Unknown | Unknown | Unknown | 100% (210) |
| DMS176 | *dma51* | C24G6.2 | *hir-1* | tyrosine | plasma | 82% (134) |
| DMS177 | *dma52* | T14B4.6 | *dpy-2* | collagen | extracellular | 92% (145) |
| DMS127 | *dma22* | EGAP7.1 | *dpy-3* | collagen | plasma | 75% (128) |
| DMS180 | *dma53* | C46A5.4 | *perl-1* | animal | extracellular | 88% (105) |

**Fig. 2.**
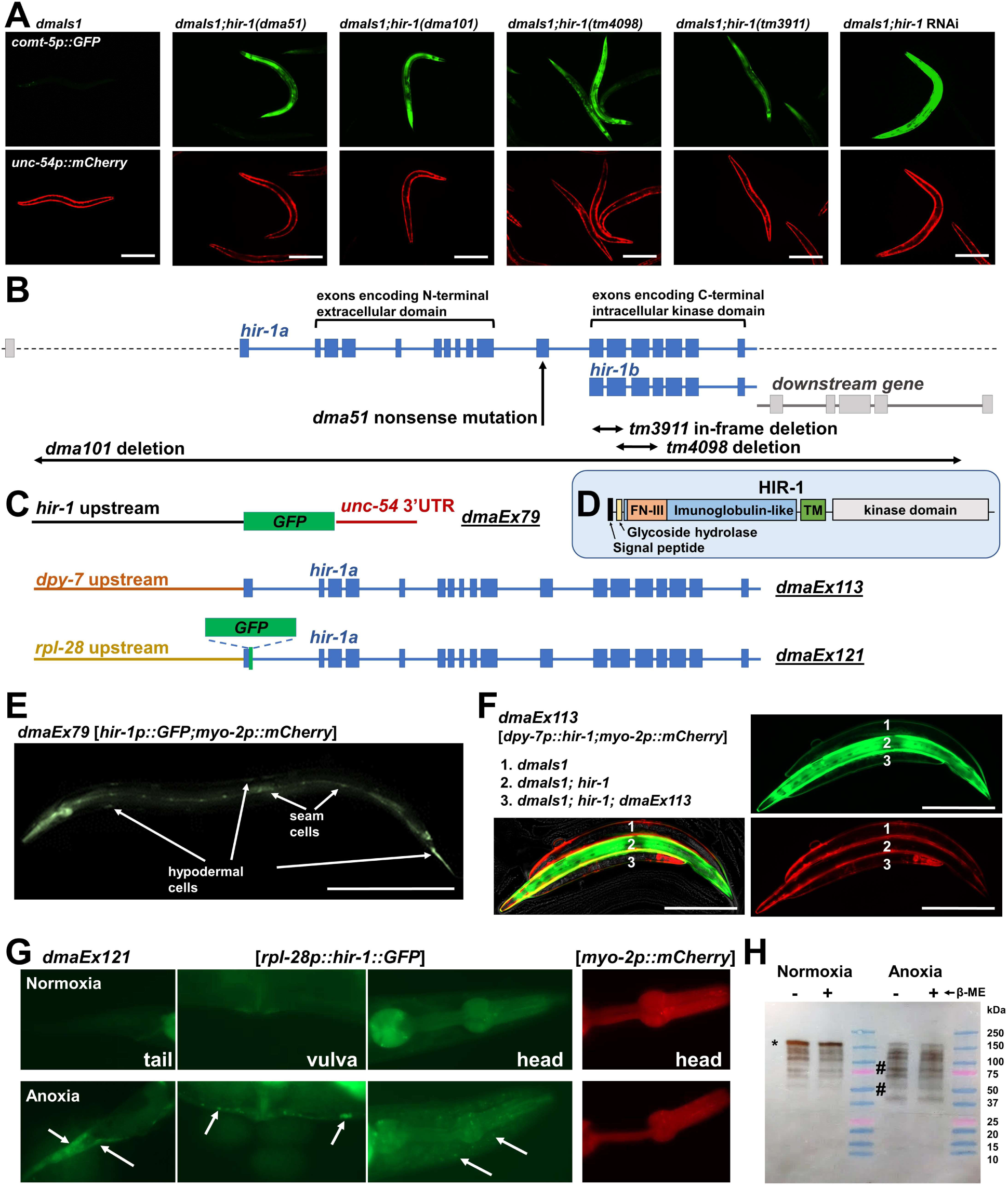
HIR-1 is a cell-autonomous regulator of signaling responses to hypoxia. **A)** Fluorescent images of transgenic animals carrying *comt*-*5p::GFP* transgene with various *hir*-*1* alleles. Representative of >100 animals. **B)** Schematic diagram of *hir*-*1* with indicated LOF alleles isolated from EMS mutagenesis (*dma51*), CRIPSR-Cas9 (*dma101*) and UV mutagenesis (*tm3911* and *tm4098*). **C)** Schematic map of various *hir*-*1* transgenes. **D)** Schematic representing domain arrangement in HIR-1. **E)** Fluorescent image of an animal carrying the transcriptional reporter *hir*-*1p::GFP*. Arrows indicate GFP signals in the hypoderm and seam cells. Representative of >100 animals. **F)** Fluorescent images of transgenic animals expressing *comt*-*5p::GFP* (worm 1) with *hir*-*1(dma101)* (worm 2) and *hir*-*1(dma101)*; *hir*-*1(*+) array *dmEx113* (worm 3); (n = 100). 23% of animals found to be rescued by *dmaEx113*. Animals carrying extrachromosomal transgene *dmaEx113 [dpy*-*7p::hir*-*1;myo*-*2::mCherry]* have mCherry fluorescence in the pharynx. **G)** Fluorescent images of transgenic animals carrying translational reporter of *hir*-*1* [*rpl*-*28p::hir*-*1sig::GFP::hir*-*1cod; myo*-*2p::mCherry]*. Arrows show aggregated GFP signal. The transgenic marker *myo*-*2p::mCherry* expressed in the pharyngeal muscles is also shown. n ≥ 10 total animals for each group with N ≥ 3 independent biological replicates. **H)** SDS-PAGE (in the presense or absence of the reducing agent β-mercaptoethanol) and western blot analysis of crude extracts of transgenic animals expressing HIR-1 ::GFP. The star indicates high-molecular weight species of GFP::HIR-1 and the hash marks indicate processed GFP::HIR-1. Representative of 3 independent experiments. Scale bars: 300 μm.

Because *hir*-*1* inactivation phenocopied exposure to hypoxia in *comt*-*5p::GFP* activation, we wondered whether hypoxia directly inhibits HIR-1. We generated a translational GFP reporter of HIR-1 (Fig. 2D) and observed decreased levels of full-length SDS-soluble HIR-1::GFP in hypoxia-treated animals (Fig. 2H). In contrast, a range of partial-length HIR-1::GFP species increased in abundance, suggesting that hypoxia leads to HIR-1 proteolytic processing (Fig. 2H). Moreover, hypoxia increased the abundance of HIR-1::GFP in the protein pool that did not migrate in the SDS-PAGE, suggesting partial insolubility of HIR-1 induced by hypoxia (Fig. S2). By direct imaging of HIR-1::GFP, we observed that hypoxia induced formation of HIR-1::GFP foci without affecting the transgenic co-injection marker *unc*-*54p::mCherry* in the pharyngeal muscles (Fig. 2G). These data indicate that hypoxia regulates HIR-1::GFP proteins and thereby control gene expression of *comt*-*5* downstream of HIR-1.

### Inactivation of genes essential for extracellular matrix integrity mimics hypoxia and the *hir*-*1(-)*-induced transcriptional response

By genetic linkage mapping and whole-genome sequencing, we identified additional genes defined by other mutants isolated from EMS screens with constitutive *comt*-*5p::GFP* expression. *dma52* is a Q4R mutation in *dpy*-*2*. *dma22* is a G162R mutation in *dpy*-*3*. *dma53* defines a previously uncharacterized gene *C46A5*.*4*, named as *perl*-*1*, which encodes a heme peroxidases-like-1 (Table 1, Fig. 3A and 3B). RNAi directed against *dpy*-*2*, *dpy*-*3* or *perl*-*1* or partial deletion alleles of these genes phenocopied EMS-derived mutations in activating constitutive expression of *comt*-*5p::GFP* under normoxia. Notably, both *dpy*-*2* and *dpy*-*3* encode cuticle collagens (30) (Fig. 3C) while *perl*-*1* is paralogous to *mlt*-*7*, which encodes a heme peroxidase required for proper crosslinking of ECM collagens (31). Both *dpy*-*3* and *dpy*-*2* are expressed in hypoderm and their LOF mutations, including EMS-derived alleles, resulted in abnormal animal morphology (dumpy phenotype) and mimicked hypoxia-induced *comt*-*5p::GFP* activation (Fig. 3A). We performed 8 independent genetic screens of over 100,000 haploid genomes and observed several *comt*-*5p::GFP* activating mutants that are unable to propagate and these mutants commonly exhibited cuticle morphological defects, such as blistering or dumpy phenotypes. We used RNAi to knock-down additional genes directly responsible for cuticle biosynthesis and observed that RNAi against cuticular collagens (*dpy*-*7*, *dpy*-*10*, *bli*-*6*), collagenase inhibitor (*bli*-*5*), another heme peroxidase (*mlt*-*7*), dual oxidase complex (*bli*-*3* and *tsp*-*15*), subtilisin-like protease (*bli*-*4*) or thioredoxin (*dpy-11*) activated *comt*-*5p::GFP* in the hypoderm (Table S1, Fig. S3A). Cuticle biosynthesis in the hypoderm is essential for molting and several cuticular genes are regulated by the molting-controlling transcription factors including HBL-1, NHR-23 and NHR-25 (32). RNAi against these transcription factor genes resulted in molting defects accompanied by activation of *comt*-*5p::GFP* (Table S1, Fig. S3A). These results indicate that impaired hypodermal ECM integrity, which results in dumpy or blistering morphological defects, activates *comt*-*5p::GFP*.

**Fig. 3.**
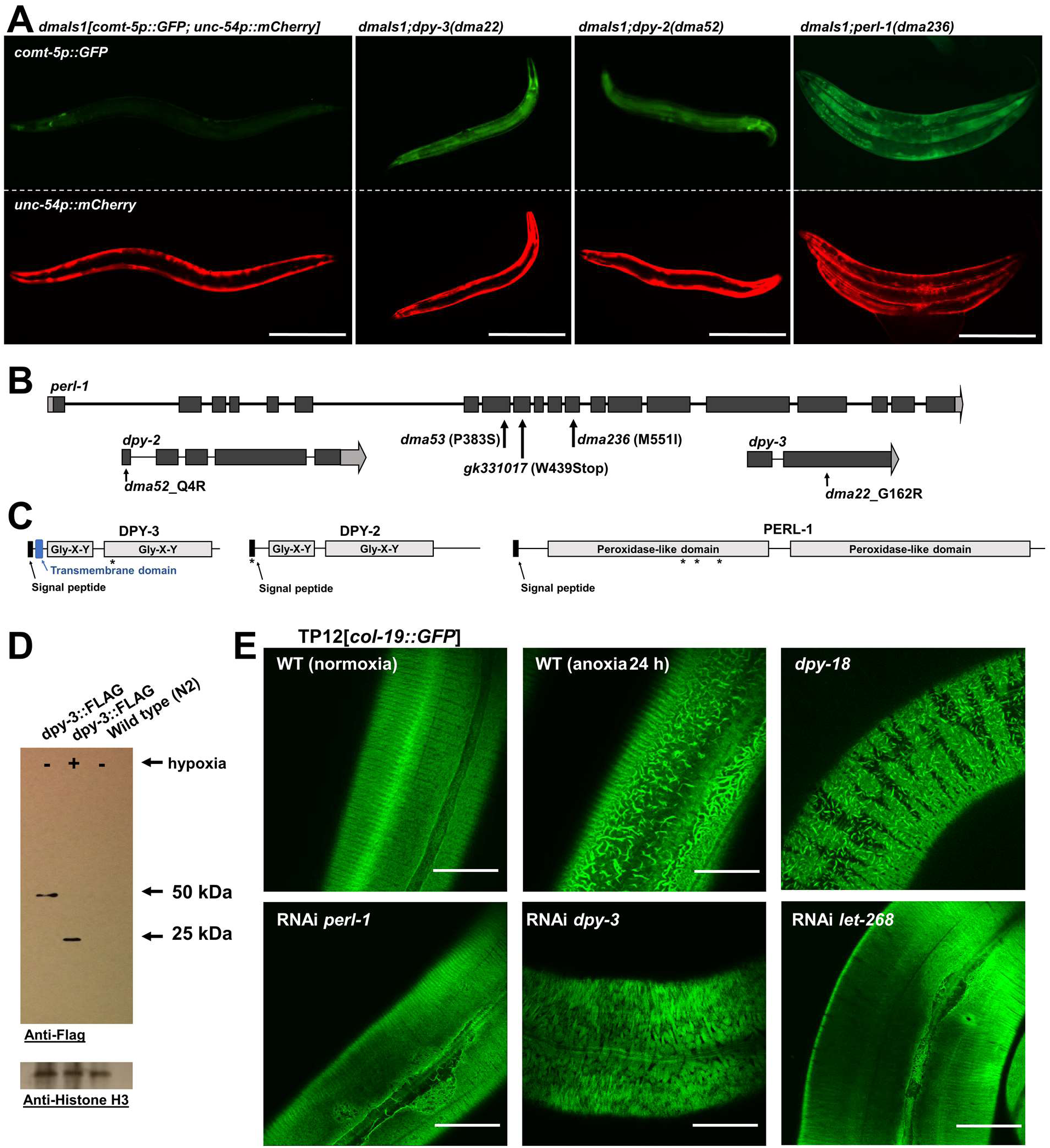
Deficiency of specific cuticular genes mimics hypoxia-induced *comt*-*5p::GFP* activation. **A)** Fluorescent images of *comt*-*5p::GFP* expression in the hypoderm of wild-type animals or mutants lacking *dpy*-*2*, *dpy*-*3* or *perl*-*1*. Representative of >100 animals. **B)** Schematic gene structures of *dpy*-*2*, *dpy*-*3* and *perl*-*1* showing the positions of EMS-derived LOF alleles. **C)** Protein domain organization of DPY-2, DPY-3 and PERL-1 with ^∗^ indicating the positions of EMS-derived mutations. The G162R mutation in DPY-3 is in the conserved collagen domain, and the Q4R mutation in DPY-2 is in the signal peptide. The missense mutations in the PERL-1 are in the N-terminal peroxidase-like domain. **D)** Western blot analysis of WT animals or animals expressing the DPY-3-FLAG translational reporter exposed to normoxia or hypoxia. Representative of 2 independent experiments. **E)** Fluorescent images of animals carrying the *col-19::GFP* translational reporter exposed to normoxia, hypoxia, or *dpy*-*18(-)*, *dpy*-*3(-)*, *perl-1(-)* and RNAi directed against *let*-*268*. Representative of 10 animals. Scale bars: 25 μm. **B)** Fluorescent images of *comt*-*5p::GFP* expression in animals treated with control or RNAi against the indicated genes essential for cuticle integrity. Representative of >100 animals. Scale bars: 300 μm.

### Hypoxic stress results in the remodeling of ECM and cuticle disintegration

Since impaired hypodermal ECM integrity increased expression of *comt*-*5p::GFP* we wondered whether hypoxia, which also leads to activation of *comt*-*5p::GFP*, leads to the remodeling of hypodermal ECM. To test whether collagens proximal to the hypodermal plasma membrane are directly affected by hypoxia, we generated transgenic animals with FLAG-tagged collagen DPY-3. By Western blot analysis we found that the molecular weight of DPY-3::FLAG was markedly altered in animals exposed to hypoxia for 24 hours (Fig. 3D), indicative of altered DPY-3::FLAG cross-linking. In contrast, another cuticle collagen COL-19 did not exhibit altered molecular weight after exposure to hypoxia or genetic conditions increasing expression of *comt*-*5p::GFP*, including RNAi against *perl*-*1* and *dpy*-*3* (Fig. S3B). Translational reporter COL-19-GFP normally localizes to the circumferential annular rings and longitudinal alae of the adult exoskeleton and genetic conditions activating *comt*-*5p::GFP* lead to disorganization of COL-19-GFP (33). We examined COL-19::GFP in WT animals exposed to hypoxia for 24 hours (nearly 0% O_2_) and observed disorganized COL-19::GFP distribution proximal to the alae (Fig. 3E).

Cuticle biosynthesis begins in the endoplasmic reticulum (ER) lumen when collagens are hydroxylated by prolyl and lysyl hydroxylases allowing its trimerization, glycosylation and secretion into ECM (34,35). Although there are four ER procollagen hydroxylases in *C*. *elegans* (*dpy*-*18*, *phy*-*2*, *phy*-*3 or phy*-*4*), which are partially genetically redundant (36), the hypomorphic allele of *dpy*-*18* caused disorganization of the COL-19::GFP, which was similar to the pattern observed in worms exposed to hypoxia. We also observed cuticle disorganization in animals with inactivated *let*-*268*, the sole *C*. *elegans* lysyl hydroxylase, and *perl*-*1* whereas inactivated *dpy*-*3* resulted in altered GFP pattern characterized by a complete loss of the cuticular furrows (Fig. 3E). Interestingly, LET-268 activity appears to be limited to collagen type IV localized in the basement membrane but not cuticular collagens (37,38). We performed RNAi against *let*-*268* and observed larval arrest with *comt*-*5p::GFP* activation (Table S1, Fig. 3A). On the other hand, the RNAi against *emb*-*9* encoding collagen type IV did not increase expression of the *comt*-*p::GFP* (Fig. 3A). The inactivation of *let*-*268* leads to accumulation of the collagen type IV in the ER (38) suggesting that inactivation of *let*-*268* activates *comt*-*5p::GFP* indirectly likely through affecting maturation of ECM proteins in the ER. These data indicate that insufficient cuticular collagen modification including hydroxylation mimics hypoxia or *hir*-*1*-induced *comt*-*5p::GFP* activation in *C*. *elegans*.

In addition, WT animals exposed to hypoxia for 24 hours (nearly 0% O_2_), *perl*-*1*, *dpy*-*3* or *dpy*-*18* mutants or *let*-*268* knockdowns by RNAi exhibited exacerbated sensitivity to osmotic stress, characteristic of mutants with disrupted cuticle integrity (39), and is likely caused by increased cuticle permeability to water (Fig. S3C). Furthermore, direct permeability assays with the cuticle impermeable dye Hoechst 22358 (40) revealed intercalation of the dye in the nuclei only in *perl*-*1* and *dpy*-*3* mutants, *let*-*268* knockdowns and hypoxia-treated WT animals but not WT animals under normoxia (Fig. S3D). These data provide multiple independent lines of evidence that cuticle in animals exposed to hypoxia exhibits altered integrity characterized by increased permeability.

### NHR-49 and MDT-15 mediate *comt*-*5* transcriptional response to hypoxia

To identify specific transcription factors that drive *comt*-*5p::GFP* expression in response to hypoxia and ECM remodeling, we sought second-site suppressor mutations of the most penetrant *comt*-*5p::GFP*-activating mutation *dma11* isolated from EMS screens (Table 1). The gene defined by *dma11* remains as yet unidentified. Nonetheless, we isolated two independent suppressing alleles *dma53* and *dma54* and used linkage analysis and RNAi phenocopying to identify them as mutations of *mdt*-*15* and *nhr*-*49*, respectively. *dma53* is a nonsense mutation leading to a premature stop codon in *mdt*-*15*, and *dma54* is a missense mutation G33R in *nhr49* (Fig. 4B). NHR-49 and MDT-15 are transcriptional regulators that physically interact to regulate lipid homeostasis (41). Protein sequence analysis of orthologous nuclear hormone receptors revealed that Gly^33^ is in the conserved DNA binding domain among all examined sequences, including the vertebrate orthologue HNF4 (Fig. 4D). To verify that LOF of *nhr*-*49* also suppressed *hir*-*1*-induced *comt*-*5p::GFP* activation, we crossed the *nhr*-*49* null allele *nr2041* with *hir*-*1* mutants and found that the *hir*-*1*; *nhr*-*49* double mutants exhibited suppressed *comt*-*5p::GFP*. Moreover, hypoxia did not activate *comt*-*5p::GFP* in *nhr*-*49* or *mdt*-*15* null mutants whereas the gain-of-function mutations *nhr*-*49(et7)* and *mdt*-*15*(*et14*) showed constitutively activated *comt*-*5p::GFP* under normoxia (Fig. 4A). Western blot and qRT-PCR analysis confirmed the requirement for NHR-49 and MDT-15 in the activation of *comt*-*5p::GFP* in *hir*-*1* mutants (Fig. 4C). qRT-PCR analysis also confirmed that both *nhr*-*49* and *mdt*-*15* were required for hypoxic induction of *comt*-*5* (Fig. 4E). These data indicate that transcriptional activation of *comt*-*5* by hypoxia or LOF of HIR-1 requires NHR-49 and MDT-15.

**Fig. 4.**
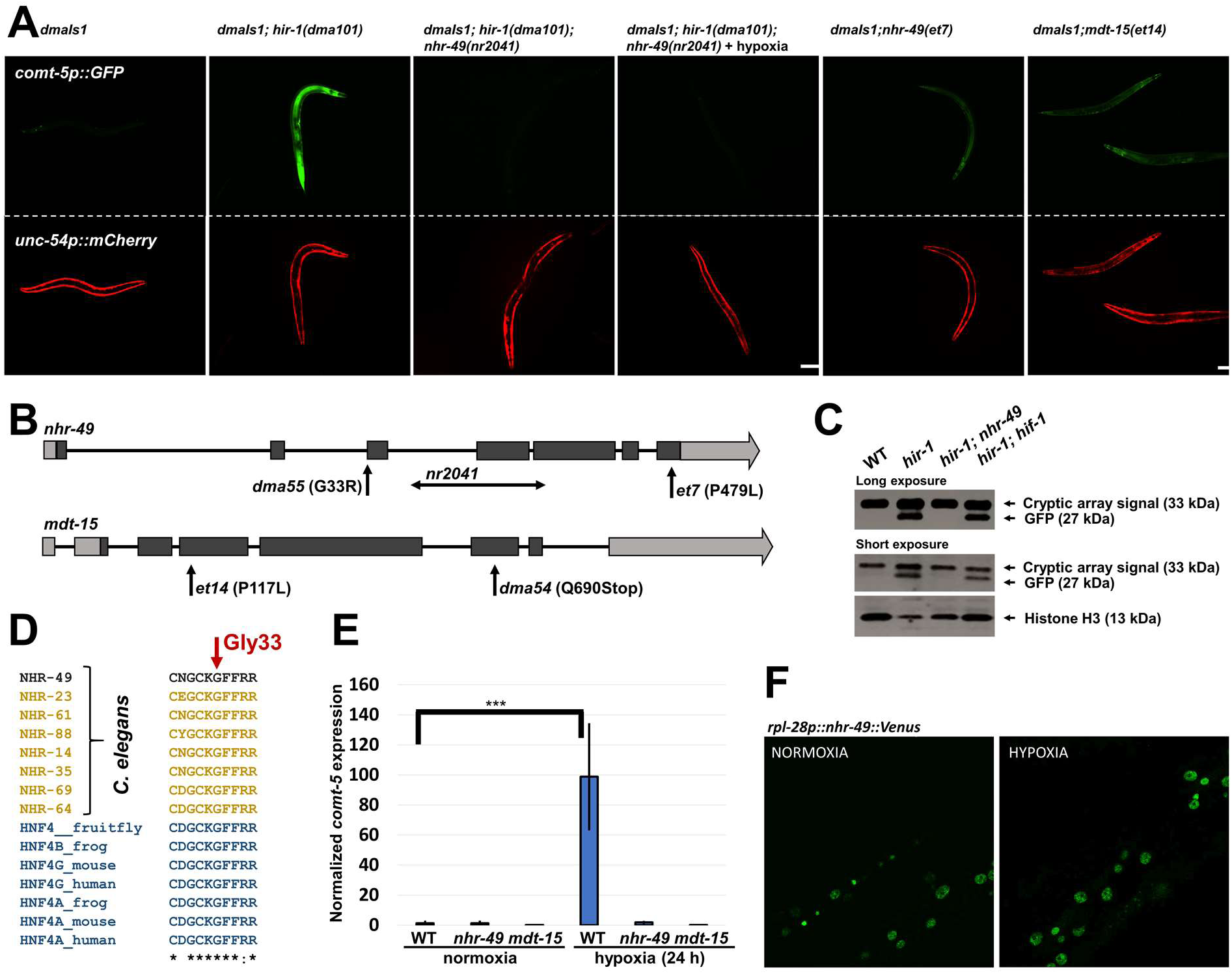
HIR-1 regulation of *comt*-*5p::GFP* requires the transcription factors NHR-49 and MDT-15. **A)** Fluorescent images of transgenic animals carrying *comt*-*5p::GFP*, and LOF and GOF of the indicated genes. Representative of >100 animals. **B)** Schematic gene structures of *nhr*-*49* and *mdt*-*15* showing the position of EMS-derived mutations. **C)** Western blot analysis of *comt*-*5p::GFP* expression in *hir*-*1*, *hir*-*1; hif*-*1*, and *hir*-*1; nhr*-*49* mutants. Representative of 2 independent experiments. **D)** Protein sequence alignment of animal proteins orthologous to NHR-49 showing that Gly^33^ in NHR-49 is fully conserved in orthologues including human hepatocyte nuclear factor 4 (HNF4). **E)** qRT-PCR analysis of endogenous *comt*-*5* expression in WT animals or *nhr*-*49* or *mdt*-*15* mutants exposed to normoxia or hypoxia. N ≥ 200 total animals for each group with 3 independent biological replicates; ^∗∗∗^ indicates P < 0.001. Scale bars: 300 μm. **F)** Fluorescent images of transgenic animals carrying *rpl*-*28p::nhr*-*49::Venus* showing hypodermal nuclei exposed to normoxia or hypoxia.

NHR-49 and MDT-15 orchestrate lipid homeostasis in *C*. *elegans* (42) and mediate response to changes in lipid metabolic cues and temperature (11,42). We generated a transgenic strain expressing *nhr*-*49::Venus* under the ubiquitous *rpl*-*28* promoter. We did not observe altered pattern of the subcellular localization of NHR-49::Venus after hypoxia, suggesting that NHR-49 is not regulated at the level of nuclear translocation. In addition, we observed that *comt*-*5p::GFP* induction by *hir*-*1* inactivation was limited to adulthood and increased with age, whereas *nhr*-*49* gain-of-function mutation or hypoxia activated *comt*-*5p::GFP* in all developmental stages (Fig. S4A). These results support that NHR-49 is essential for HIR-1-dependent transcriptional response but plays broader roles than HIR-1.

### Mechanisms of HIR-1 regulation by hypoxia-induced ECM remodeling

The genetic evidence connecting hypoxia, ECM integrity and *hir*-*1* suggests that exposure to hypoxia induces remodeling of the ECM, leading to inhibition of HIR-1 and subsequent activation of *comt*-*5p::GFP*. The intracellular domain of HIR-1 is structurally similar to the proto-oncogene receptor RET and fibronectin growth factor (FGF) receptors, with conserved catalytic sites essential for autophosphorylation (Fig. 5A). We wondered whether the HIR-1 kinase activity could be inhibited by depletion of ATP due to hypoxia. We exposed animals to rotenone, an inhibitor of the oxidative phosphorylation, but did not observe increased expression of *comt*-*5p::GFP*. The extracellular domain of HIR-1 resembles immunoglobulin-like fold (a.a. 80-446) including fibronectin type III-like fold (a.a. 84-170), domains common in many receptors previously identified as interacting with ECM proteins (43,44). FGF4 is a ligand implicated in mediating ECM sensing to regulate trophoblast stem cell fate (13). Inactivation of *C*. *elegans* FGF-encoding gene *let-756* by RNAi led to constitutive activation of *comt*-*5p::GFP* under normoxia, phenocopying *hir*-*1* LOF mutants (Fig. 5B). We generated an HA-tagged translational reporter of *let*-*756* and observed hypoxia-induced change in LET-756 molecular weights (Fig. 5C). These findings indicate co-regulation of HIR-1 and LET-756, supporting the notion that HIR-1 acts with FGF-like proteins to sense ECM remodeling upon hypoxia, which would attenuate binding of FGF-like ligands and thereby inactivating HIR-1.

**Fig. 5.**
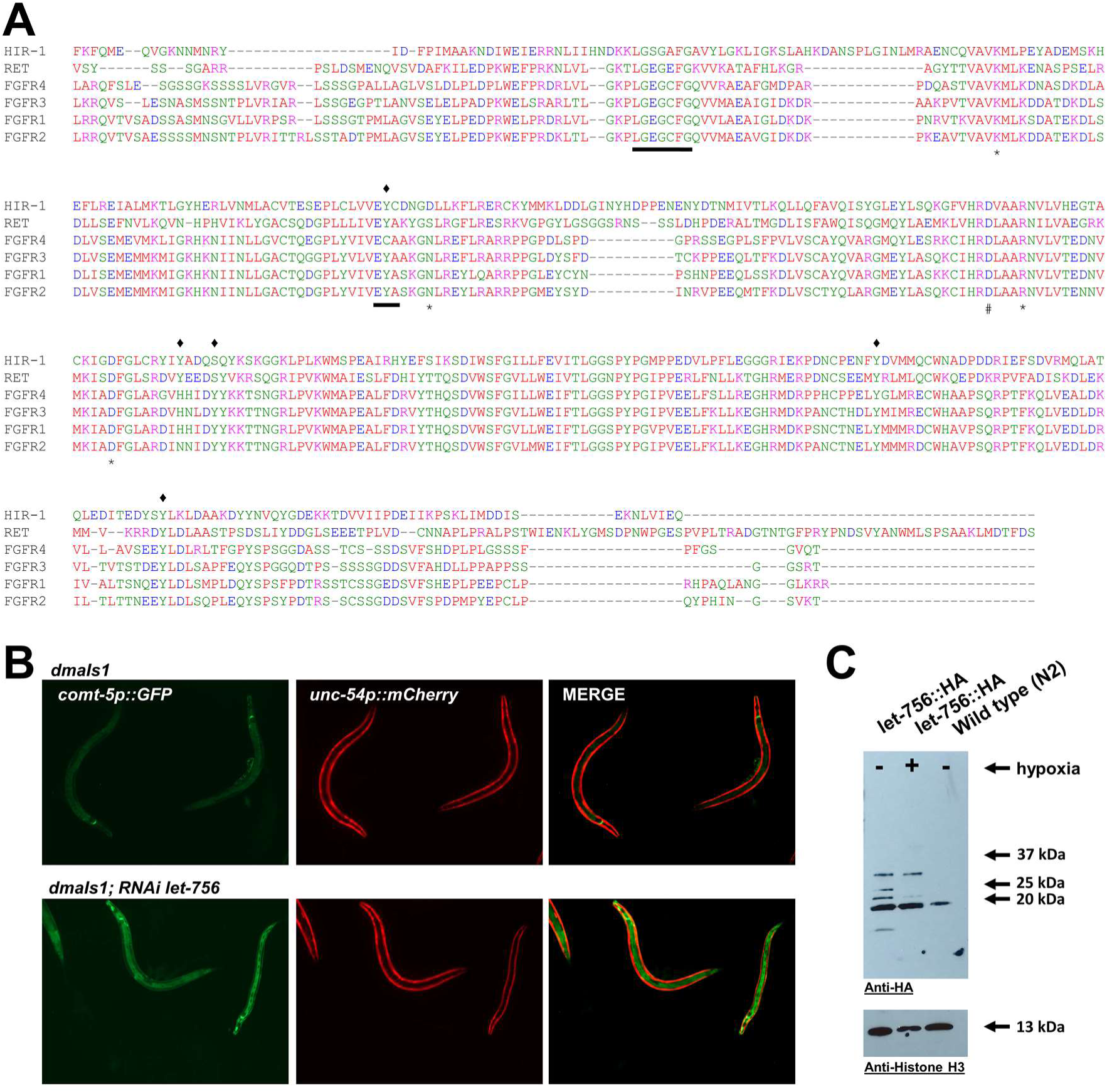
LET-756 is potential lignaf for HIR-1. **A)** Protein sequence alignment of intracellular kinase domains in human RTKs orthologous to HIR-1 indicating that HIR-1 possesses a conserved ATP binding site designated by ^∗^, a substrate proton acceptor site designated by ‐, nucleotide binding sites designated by lines underlining the sequences, and conserved autophosphorylated tyrosine/serine residues found to be modified in human orthologues are designated by ♦ above the sequences. **B)** Fluorescent images of *comt*-*5p::GFP* animals showing GFP induction elicited by RNAi against *let*-*756*.

### HIR-1 regulates a broad genetic program controlling ECM homeostasis and protects against severe hypoxic stress

To systematically identify genes differentially regulated by *hir*-*1* in relation to hypoxia, we performed RNA-seq analysis of *hir*-*1* mutants and compared transcriptomes of WT animals under normoxia or hypoxia and *hir*-*1* mutants (Fig. S1B, S5A and S5B). Expression of genes that regulate *comt*-*5p::GFP*, including *dpy*-*2*, *dpy*-*3*, *perl*-*1*, was increased by both hypoxia and *hir*-*1* inactivation (Fig. S2A). We used qRT-PCR to quantify the expression levels of genes encoding collagens, procollagen hydroxylases, peroxidases and transcription factors mediating molting processes in *hir*-*1*, *hir*-*1*; *nhr*-*49* and *hir*-*1*; *hif*-*1* mutants. All examined genes showed increased expression in *hir*-*1* and *hir*-*1*; *hif*-*1* mutants and decreased expression in *hir*-*1*; *nhr*-*49* mutants (Fig. S2B). Furthermore, we found that genes involved in molting and cuticle integrity showed increased expression in *hir*-*1* mutants, including members of collagen-encoding *dpy* genes whose inactivation can cause the Dumpy phenotype (Data File S1). Several collagen-encoding non-Dumpy genes, including *col*-*17* and *col*-*41*, were suppressed (Fig. S5B). Collagens are the main structural components of the ECM and cuticle, and the epicuticle contains lipids that regulate its permeability (46). We found that *hir*-*1* mutants exhibited increased expression of genes involved in lipid metabolism, including *acs-2*, a target gene of NHR-49 and key regulator of fatty acid homeostasis (42) (Fig. S5B). LOF of *hir*-*1* led to disrupted cuticle integrity, supporting that *hir*-*1* mutants exhibit more permeable cuticles (Fig. 6B and 6C, and S3F). The cuticle defects of *hir*-*1* mutants was not fully rescued by *nhr*-*49* LOF, indicating that defects in ECM integrity of *hir*-*1* mutants under normoxia involves abnormal activation of additional unidentified transcription factors (Fig. S4B).

**Fig. 6.**
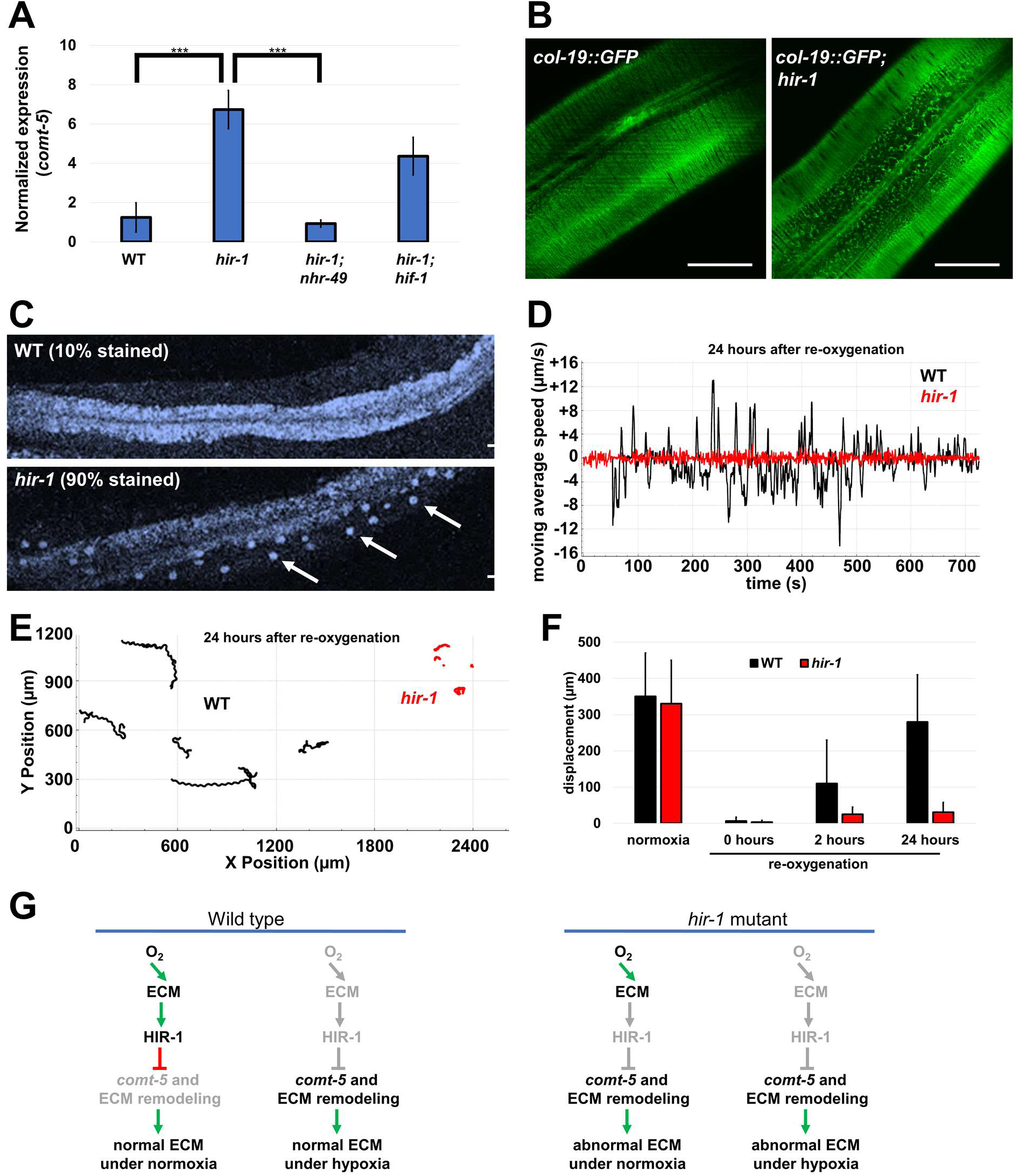
HIR-1 mediates ECM homeostasis and anoxia resistance. **A)** qRT-PCR analysis of *comt*-*5* expression in WT, *hir*-*1*(*dma101*), *hir*-*1<(dma101); hif*-*1(ia04)* and *hir*-*1 (dma101); nhr*-*49(nr2041)* mutant animals. N ≥ 200 total animals for each group with 3 independent biological replicates; ^∗∗∗^ indicates P < 0.001. **B)** Fluorescent images of animals expressing the *col*-*19::GFP* translational reporter in the presence or absence of *hir*-*1* RNAi. Representative of 10 animals. **C)** Cuticle integrity as detected by nuclear staining with Hoechst 22358 in the hypoderm of WT or *hir*-*1(dma101)* animals. Numbers represent percentages of animals with stained nuclei (n = 10 animals). **D)** Representative plot showing the average locomotion speed of a WT or *hir*-*1* worm in a 10-minute time frame recorded after 24 hours of re-oxygenation. **E)** Plot showing representative locomotor tracks of WT and *hir*-*1* animals in a 10-minute time frame recorded after 24 hours of re-oxygenation. Position is detected based on the mid-point of the worm. **F)** Displacement of WT and *hir*-*1* mutants in a 10-minute time frame recorded before exposure to hypoxia, and after 0 - 10 min, 2 and 24 hours of re-oxygenation; ^∗∗∗^ indicates P < 0.001. N=10 animals for each group with 3 independent biological replicates. **G)** Schematic model of the proposed HIR-1 pathway that mediates cellular response to hypoxia-induced ECM remodeling to maintain ECM homeostasis in *C*. *elegans*. Inactive components are shown in gray and activated components are shown in black. Scale bars: 25 μm.

We next tested whether the genetic program mediated by HIR-1 signaling pathway was essential for resistance to severe hypoxic stress. We exposed 1-day old WT and *hir*-*1 (dma101)* mutants to anoxia for 40 hours and subsequently compared their locomotion behavior after re-oxygenation. All tested animals were in suspended animation-like state unable to respond to external stimuli (mechanic touch or UV light) during anoxia and immediately after re-oxygenation, which was followed by behavioral recovery in crawling (defined as displacement) and pharyngeal pumping. We found that the displacement of re-oxygenated WT animals gradually recovered, whereas *hir*-*1* mutants had markedly severe locomotion defects that did not improve even after 24 hours of re-oxygenation and culminated in enhanced rates of organismic death (Fig. 6D-F). These results indicate that HIR-1 plays key roles to control a broad genetic program that guards against maladaptive cuticle collagen and ECM homeostasis for animal survival upon exposure to severe hypoxia (Fig. 6G).

## DISCUSSION

How cells in multicellular organisms sense hypoxia-remodeled ECM to promote cuticle homeostasis and facilitate animal adaptation to severe hypoxia is unknown. From genetic screens, we identified HIR-1 as a key mediator of the transcriptional response to hypoxia-induced ECM remodeling (47,48). Altered expression pattern of GFP-tagged HIR-1 in animals exposed to hypoxia suggests that HIR-1 is directly regulated by hypoxia-induced ECM remodeling. The observed HIR-1::GFP foci are reminiscent of aggregation and proteolytic processing, a previously observed effect of hypoxia on other proteins in *C*. *elegans* (27,28). We do not exclude a possibility of receptor clustering, internalization or block of the membrane trafficking that would affect activity of HIR-1 or its interacting partners. HIR-1 does not possess the discoidin domain, which is responsible for interaction with collagens to sense ECM remodeling (45). Our genetic evidence suggests that LET-756 homologous to mammalian FGFs is a likely HIR-1 ligand, whose binding to HIR-1 might be modulated by hypoxia-induced ECM remodeling. How precisely HIR-1 is regulated by ECM remodeling in coordination with its ligand and, in turn, transduces intracellular signaling for ECM homeostasis awaits further studies.

The integrity and composition of ECM are sensitive to oxygen availability due to oxygen-dependent enzymatic activities of prolyl and lysyl hydroxylases in the ER as well as dual oxidases in the extracellular space (31,49). We found that procollagen hydroxylase or dual oxidase inactivation by RNAi phenocopied hypoxia-induced activation of *comt*-*5p::GFP*. LOF phenotypes of these oxygen-dependent enzymes suggest that they likely mediate hypoxic response directly, whereas their HIF-induced increase in expression might constitute a homeostatic response to eventually restore their activity in the cell. Depending on the tissue-specific range of oxygen levels to which these enzymes are sensitive, procollagen hydroxylases and extracellular oxidases likely act as cellular oxygen sensors that mediate responses to varying degrees of hypoxia. We propose that such oxygen sensing in the ER or extracellular space acts in parallel to cytosolic EGLN oxygen sensors to mediate HIF-independent transcriptional programs to induce ECM remodeling and inhibit HIR-1 to maintain ECM homeostasis and animal survival under severe hypoxic conditions.

Hypoxia-induced ECM remodeling occurs in cultured tumor cells (5) and ECM surrounding bone marrow tumor cells exhibit increased stiffness that triggers metastasis (50). Increased stiffness of the ECM due to enhanced crosslinking of ECM proteins is mediated by increased expression of lysyl hydroxylase and lysyl oxidase and increased collagen deposition (5,51,52). Similarly, we also found hypoxia increased the expression of genes encoding procollagen lysyl oxidase, ER hydroxylases, and collagens in *C*. *elegans*, indicating conserved transcriptional responses that promote homeostatic ECM remodeling, which may have been coopted by tumors to facilitate hypoxic survival and metastasis. ECM remodeling during aging through differential regulation of collagen-encoding genes contributes to extension of *C*. *elegans* longevity (53), consistent with the notion that homeostatic ECM remodeling mediated by the HIR-1 pathway promotes cellular and organismic resistance to hypoxic stresses.

We found that proper HIR-1 regulation is important for *C*. *elegans* adaptation to hypoxic stress and re-oxygenation while its downstream transcriptional effectors include not only ECM components but also a non-ECM gene *comt*-*5*. *comt*-*5* is predicted to encode a catecholamine-degrading enzyme, and its upregulation by hypoxia likely help decrease catecholamine levels to alleviate the toxicity of their oxidized forms under hypoxia, while its constitutive expression in *hir*-*1* mutants under normoxia is likely detrimental. Dopamine targets peripheral tissues to maintain xenobiotic stress resistance in *C*. *elegans* (54), however, whether upregulation of *comt*-*5* is required for proper resistance to hypoxic and/or xenobiotic stress awaits further studies. Although we used primarily *comt*-*5*::GFP reporters for gene and pathway discoveries, transcriptomic analysis revealed that HIR-1 regulates a broad genetic program responsible for remodeling of ECM, altering cuticle integrity and reprograming lipid metabolism.

Our findings suggest that *hir*-*1*- and *nhr*-*49*-mediated reprogramming of lipid metabolism likely also contributes to hypoxic survival in addition to ECM homeostasis. Hypoxic regulation of genes involved in cuticular lipid synthesis can promote cuticle permeability and adaptation to hypoxia in *Arabidopsis thaliana* (55). Lipid metabolic reprogramming also contributes to tumor development and hypoxic resistance in mammals (56). Upstream of NHR-49, the regulatory axis from oxygen, ER hydroxylases, and collagen to RTKs appears to share features common in both *C*. *elegans* and humans. We thus propose that the HIR-1 pathway is evolutionarily conserved to mediate cellular response to hypoxia and ECM remodeling in diverse organisms. Numerous human RTKs, including FGFRs, have been implicated in driving tumor survival and metastasis (57,58), and ECM remodeling is essential for the progression of cancer, tissue fibrosis and many other diseases involving ECM dysregulation (59). Given the central role of HIR-1 in mediating cellular response through ECM remodeling to promote resistance to hypoxia, the human counterpart of HIR-1 may, once verified, be a promising therapeutic target for solid tumors that survive severe hypoxia and metastasize through ECM remodeling.

## MATERIALS AND METHODS

### *C*. *elegans* strains

Animals were maintained under standard procedure with nematode growth media (NGM) plates unless otherwise stated. Bristol strain N2 was used as wild type and Hawaiian strain CB4856 was used for the linkage analysis of the mutants (60,61). *hir*-*1* null alleles were generated by CRISPR to induce double stranded breaks and subsequent non-homologous end joining caused a deletion of *hir*-*1*. Feeding RNAi was performed as previously described (62). Transgenic strains were generated by germline transformation as described (63). Transgenic constructs were co-injected with dominant *unc*-*54p*::mCherry or *myo*-*2*::mCherry markers, and stable extrachromosomal lines of mCherry+ animals were established. Transgenic strains used were DMS24 *(dmaIs1*), DMS23 (*dmaIs1;hif*-*1(ia04)*), DMS22 (*dmaIs1;egl*-*9(sa307)*), DMS176 (*dmaIs1;hir*-*1(dma51)*), DMS290 (*dmaIs1;hir*-*1(dma101)*), DMS492 (*dmaIs1;hir*-*1(tm3911)*), DMS311 (*dmaIs1;hir*-*1(tm4098)*), DMS634 (*dmaIs1;hir*-*1(dma101);nhr*-*49(nr2041)*), DMS290 (*dmaIs1;hir*-*1(dma101)*, DMS99 (*dmaIs1;dma11*), DMS181(*dmaIs1;dma11;mdt*-*15(dma54)*), DMS182 (*dmaIs1;dma11;nhr*-*49(dma55)*), DMS283 (*dmaIs1;mdt*-*15(et14)*), DMS127 (*dmaIs1;dpy*-*3(dma22)*), DMS177 (*dmaIs1;dpy*-*2(dma52)*), DMS180 (*dmaIs1;perl*-*1(dma53)*), DMS582 (*qyIs44[emb*-*9::mCherry]*), DMS696 (*dmaIs1;perl*-*1(dma236)*), DMS636 (*dmaIs1; hir*-*1(dma101); dmaEx113[dpy*-*7::hir-1(+); myo*-*2::mCherry]*), DMS665 (*dmaEx121[rpl*-*28p::hir*-*1::SignalPeptide::gfp::hir*-*1; myo*-*2p::mCherry]*), TP12 (*kaIs12[col*-*19::GFP]*).

### Genetic screens

Stereo-epifluorescence dissecting microscope (Nikon SMZ18) was used to isolate mutants with constitutive expression of *comt*-*5p::GFP* reporters after ethyl methanesulfonate (EMS)-induced mutagenesis, as described previously (2,11). Mutants were mapped genetically by single nucleotide polymorphisms-based linkage analysis using the Hawaiian *C*. *elegans* strain CB4856 and then were sequenced by whole-genome sequencing to obtain lists of candidate genes. Genes with putative causal mutations were verified by RNA interference that can phenocopy mutants and the causality of mutation was subsequently confirmed by transformation rescue of mutants with wild-type alleles as transgenes.

### Environmental stress assays

Hypoxia chamber with ProOx110 oxygen controller (Biospherix) was used for hypoxia stress with 0.5%-21% oxygen concentrations. Hypoxia incubator chamber (Applied StemCell) with constant nitrogen flow delivering was used for achieving severe hypoxia with nearly 0% oxygen (anoxia stress). In hydrogen peroxide stress assay, animals were placed on the plate with 10 mM peroxide in the NGM and observed in the 1-48-hour interval for GFP activation. For determining effects of HIF-activating compounds on *comt*-*5p::GFP*, 5 mM CoCl_2_, 5 mM KCN containing NGM plates were used. For assaying hydrogen sulfide, 0.1 mg of NaHS powder was placed onto NGM agar plate (10 ml of agar) with sealed lid by parafilm to prevent leaking of the released H_2_S gas. Animals were subsequently screened for the GFP induction in the 1-48-hour interval, and for the qRT-PCR analysis collected after 2 hours of exposure.

### Western Blot analysis

Animals were lysed in the Laemmli sample buffer (BioRad) with reducing agent beta-mercaptoethanol followed by boiling the samples for 10 minutes. The worm lysates were separated by SDS-PAGE and subsequently detected by GFP Goat polyclonal antibody (Fisher scientific – AF424) with Histone H3 antibody (AbCam - ab1791) as a loading control. For mCherry-tagged EMB-9, we used mCherry antibody 16D7 (Thermo Fisher Scientific M11217).

### Sample and library preparation for RNA sequencing

All *C*. *elegans* strains (N2(WT), *egl*-*9(sa307)*, *egl*-*9(sa307); hif*-*1(ia04)*, *hir*-*1(tm4098)*), were maintained at 20°C prior RNA extraction. For anoxia stress, we placed N2 animals into hypoxia incubator chamber (Applied StemCell) with constant nitrogen delivering for 2 hours before lysis. 1 μg of total RNA from each sample was purified by RNeasy Mini Kit from Qiagen and used for sequencing library construction. The NEBNext^®^ rRNA Depletion Kit, Agencourt RNAClean XP Beads from Beckman Coulter, NEBNext^®^ Ultra^™^ Directional RNA Library Prep Kit for Illumina^®^, Agencourt AMPure XP from Beckman Coulter and NEBNext Multiplex Oligos for Illumina were used for preparation of sequencing libraries, per manufacturers’ instruction. The Q5 Hot Start HiFi PCR Master Mix was used for PCR enrichment of the adaptor-ligated DNA. The libraries were submitted to 100 bp paired-end high throughput sequencing using Hiseq-3000 by the Center for Advanced Technology of the University of California, San Francisco.

### RNA-seq data analysis

The prinseq-lite software (0.20.4) was used (64) to trim and filter raw reads. Reads longer than 30 bp together with minimum quality score >15 were used for subsequent analyses. The Pairfq script was used for separation of paired and single reads. Clean reads were mapped to the *C*. *elegans* genome using Hisat2 (2.0.5) (65) with default parameters. The number of mapped reads were counted by featureCounts (1.5.0) (66). Differential gene expression analysis was performed using the DESeq2 package (67). Adjusted P-value ≤ 0.05 was used as the threshold to identify the differentially expressed genes. Gene ontology and KEGG pathway enrichment analyses for the differentially expressed genes were conducted using the Cytoscape plugins BiNGO (68) and ClueGO (69), respectively. Plots for the mapped reads were generated by IGVtools (70).

### Quantitative RT-PCR

Total RNAs were isolated from animals of mixed stages, with 50 μl pellet animals (>200 animals) resuspended in 250 μl lysis buffer of Quick-RNA MiniPrep kit (Zymo Research, R1055) and subsequently lysed by TissueRuptor (Motor unit “8” for 1 min). Total RNAs were extracted following instruction (Zymo Research, R1055). 2 μg RNA/sample was reverse transcribed into cDNA (BioTools, B24408). Real-time PCR was performed by using Roche LightCycler^®^96 and SYBR Green (Thermo Fisher Scientific, FERK1081) as a dsDNA-specific binding dye. qRT-PCR condition was set to 95°C for denaturation, followed by 45 cycles of 10s at 95°C, 10s at 60°C, and 20s at 72°C. Melting curve analysis was performed after the final cycle to examine the specificity of primers in each reaction. Gene expression changes were calculated by ΔΔCT method with *act*-*3* used as reference gene. Primer sequences are in table S2.

### Imaging

Animals were mounted onto 2% agarose pad with 10 mM sodium azide and imaged with EVOS FL auto digital microscope for epifluorescence imaging or the confocal Leice SPE microscope for high-resolution *col*-*19*::GFP confocal imaging. At least 3 biological replicates (≥10 animals for each replicate) were used for quantification of the designated phenotype.

### Hoechst staining

Animals were placed into liquid drops containing 2 μg/ml of the Hoechst 22358 dye diluted in M9 buffer for 15 minutes. Then the animals were picked into fresh M9 drops and subsequently placed onto 2% agarose pad with 10 mM sodium azide for imaging by the confocal Leica SPE microscope. At least 3 biological replicates (≥10 animals for each group) were used for quantification of stained animals.

### Osmotic shock assay

5-day old adult hermaphrodites were picked and placed into 200 μl liquid drop of PCR-grade distilled water drop on plastic lid of Petri plate. Time when cuticle bursted (release of insides) was monitored by eye in 15-second intervals. 3 biological replicates (10 animals without bacteria per assay) were used for statistical analysis.

### Anoxia sensitivity assay

Animals were grown at 25°C for two continuous generations in non-starving non-stressed conditions. 1-day old adult hermaphrodites were placed on the NGM plates into the anoxia chamber for 40 hours. Animals were subsequently screened for their paralysis/movement in the indicated time intervals. WormLab system (MBF Bioscience) was used for quantification of the displacement, moving average speed and tracking based on the mid-point position. At least 3 biological replicates (10 animals per assay) were used for statistical analysis.

### Statistical analysis

Data are presented as means ± S.D. unless otherwise specified with p values calculated by unpaired Student’s t-tests and one-way ANOVA.

## SUPPLEMENTARY MATERIALS

**Fig.S1. Venn diagrams for differentially regulated genes**.

**Fig.S2. RNA-seq and qRT-PCR analysis**.

**Fig.S3. ECM and cuticle integrity assays**.

**Fig.S4. HIR-1 mediates ECM homeostasis**.

**Fig.S5. Volcano plot and Heat map**

**Table S1. Genes that regulate *comt*-*5p::GFP* expression**.

**Table S2. Primer sequences**

**Data File S1. Gene ontology and KEGG pathway analyses of genes differentially regulated in *hir*-*1* mutants compared with wild-type animals**.

## ACKNOWLEDGMENTS

We thank the *Caenorhabditis* Genetics Center, National BioResource Project in Japan and the Million Mutation Project for *C*. *elegans* strains. We thank Bingying Wang and Eric Chuang for technical assistance, and Masako Asahina and Neel Singhal for discussion.

## AUTHOR CONTRIBUTIONS

R.V. and D.M. designed, performed and analyzed the experiments and wrote the manuscript. Y.L. performed RNA sequencing and bioinformatic analysis. D.M. supervised the project.

## DATA AND MATERIALS AVAILABILITY

The RNA-seq data have been deposited to X. All other data needed to evaluate the conclusions in the paper are present in the paper or the Supplementary Materials.

